# Local sleep-like cortical reactivity in the awake brain after focal injury

**DOI:** 10.1101/2019.12.19.882290

**Authors:** Simone Sarasso, Sasha D’Ambrosio, Matteo Fecchio, Silvia Casarotto, Alessandro Viganò, Cristina Landi, Giulia Mattavelli, Olivia Gosseries, Matteo Quarenghi, Steven Laureys, Guya Devalle, Mario Rosanova, Marcello Massimini

## Abstract

The functional consequences of brain injury are known to depend on neuronal alterations extending beyond the area of structural damage. Although a lateralized EEG slowing over the injured hemisphere was known since the early days of clinical neurophysiology, its electrophysiological mechanisms were not systematically investigated. In parallel, basic sleep research has thoroughly characterized the neuronal events underlying EEG slow waves in physiological conditions. These EEG events reflect brief interruptions of neuronal firing (OFF-periods) that can occur locally and have prominent consequences on network and behavioral functions. Notably, the EEG slow waves observed following focal brain injury have been never explicitly connected to local sleep-like neuronal events. In previous works, probing cortical circuits with transcranial magnetic stimulation coupled with EEG (TMS/EEG) proved as an effective way to reveal the tendency of cortical circuits to transiently plunge into silent OFF-periods. Here, using this approach, we show that the intact cortex surrounding focal brain injuries engages locally in pathological sleep-like dynamics. Specifically, we employed TMS/EEG in a cohort of thirty conscious awake patients with chronic focal and multifocal brain injuries of various etiologies. TMS systematically evoked prominent slow waves associated with sleep-like OFF-periods in the area surrounding focal cortico-subcortical lesions. These events were associated with a local disruption of signal complexity and were absent when stimulating the contralateral hemisphere. Perilesional sleep-like OFF-periods may represent a valid read-out of the electrophysiological state of discrete cortical circuits following brain injury as well as a potential target of interventions aimed at fostering functional recovery.

**One Sentence Summary:** Focal cortical injuries are associated with local intrusion of sleep-like dynamics over the perilesional areas which disrupt local signal complexity and coexist with typical wakefulness cortical reactivity patterns within the same brain.

## Introduction

The functional consequences of a focal brain lesion are due to direct structural damage as well as to alteration in the dynamics of intact connected areas*(1–5)*. Detecting the electrophysiological changes occurring in these areas and understanding their neuronal underpinnings has been so far elusive.

A lateralized slowing of the wake EEG over the area surrounding focal lesions is a classic notion derived from early EEG recordings in acute stroke patients*(6)*. However, since the use of EEG in stroke research was superseded by imaging techniques (both structural and metabolic) offering higher spatial resolution, our understanding of stroke electrophysiology in humans proceeded at a rather low pace. By contrast, over the last twenty years the cellular/network mechanism of physiological slow EEG oscillations has been clearly identified in *in vivo*, *in vitro* as well as *in silico* models*(7–9)*.

Importantly, we have learned that, during sleep, slow waves in the scalp EEG reflect the occurrence of brief interruptions of the neuronal firing (OFF-periods) associated with a slow oscillation of the membrane potential of cortical neurons*(7)*. OFF-periods are due to the tendency of cortical neurons to fall into a silent hyperpolarized state after an initial activation (a phenomenon known as cortical bistability*(7, 10)*) represents the most dramatic change in the functional regime of neuronal cells. The mechanisms for such cellular behavior rely on both intrinsic as well as network properties, including increased activity of leak and adaptation K+ channels and/or by increased inhibition within cortical circuits*(10)*. Similar alterations may also occur as a consequence of brain injury due to a disconnection of ascending activating fibers and/or a reduction of lateral excitatory connectivity*(11, 12)*.

Surprisingly, in spite of their common features and possible mechanistic determinants, the lateralized slow waves occurring after focal brain injury have never been explicitly connected to the electrophysiology of sleep slow waves.

Such a putative connection is particularly interesting when considering that slow waves and neuronal OFF-periods can occur locally not only during sleep*(13)*, but also during wakefulness (some brain regions can be silent while others are active*(14–17)*) with important consequences on behavior, including motor impairments as shown in awake, sleep-deprived rats performing a pellet reaching task*(15)*, as well as cognitive lapses in awake humans*(18)*.

In light of these evidence, we here test the hypothesis that the electrophysiological alteration affecting structurally intact perilesional areas reflects a tendency of cortical neurons to fall into a silent OFF-period, i.e. a pathological form of local sleep patterns in the awake injured brain.

OFF-periods during non-Rapid Eye Movement (NREM) sleep can be readily detected as a suppression of high frequency power associated with a slow wave*(19)* as shown in intracranial recordings*(20, 21)*. By virtue of their activity-dependent nature, cortical bistability and the associated OFF-periods can be better revealed above and beyond spontaneous activity by recording the cortical response to direct perturbations*(22–24)*. Such perturbational approach has also been used to demonstrate that silent OFF-periods are in a key position to disrupt cortico-cortical interactions*(22)*. Most important, a follow-up study showed that the same events can be detected non-invasively by applying transcranial magnetic stimulation (TMS) combined with EEG (TMS/EEG). Specifically, multisite TMS in healthy sleeping individuals as well as in unresponsive wakefulness syndrome (UWS) patients showed the ubiquitous occurrence of OFF-periods leading to a global impairment of causality and brain complexity*(25)*.

In the present work, we apply TMS/EEG to a cohort of thirty conscious awake patients with chronic focal and multifocal brain injuries. We show that OFF-periods characterize the electrophysiological state of the perilesional area surrounding a focal cortical lesion and that this alteration is associated with a disruption of local signal complexity with respect to a control stimulation site. This finding is relevant since it connects the notion of local sleep, thoroughly described in the sleep literature, to the pathophysiology of focal brain injury and stroke. Perilesional sleep-like OFF-periods may represent a valid read-out of the state of discrete cortical circuits following brain injury as well as a potential target for the development of novel therapeutic interventions and physical rehabilitation aimed at fostering functional recovery.

## Results

In this work, we assessed the EEG responses to TMS in three groups of conscious awake brain-injured patients (total n=30) of various etiologies and disease severity. Specifically, we included i) a group of ten patients affected by unilateral cortico-subcortical lesions due to an ischemic occlusion of the middle cerebral artery (MCA ischemia group; 4 F; 68± 4 years old), ii) a group of ten patients affected by severe ischemic, haemorrhagic and traumatic multifocal cortico-subcortical lesions (severe multifocal lesions group; 4 F; 53± 6.3 years old), and iii) a group of ten patients affected by unilateral lacunar ischemic or typical haemorrhagic purely subcortical lesions (subcortical lesions group; 6 F; 72± 2.6 years old). For a complete description of patient demographics and details, see Table 1.

**Table 1.**
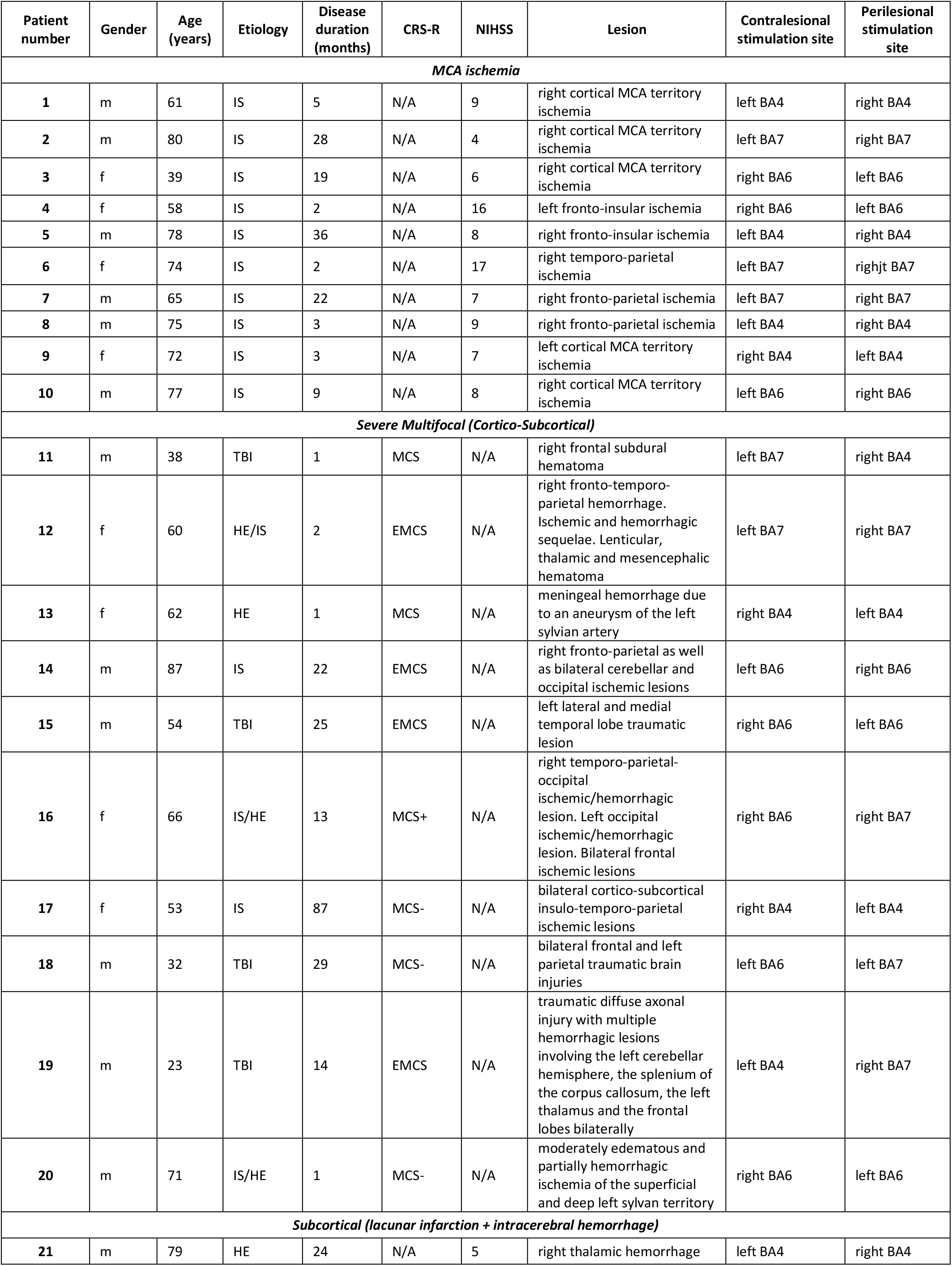
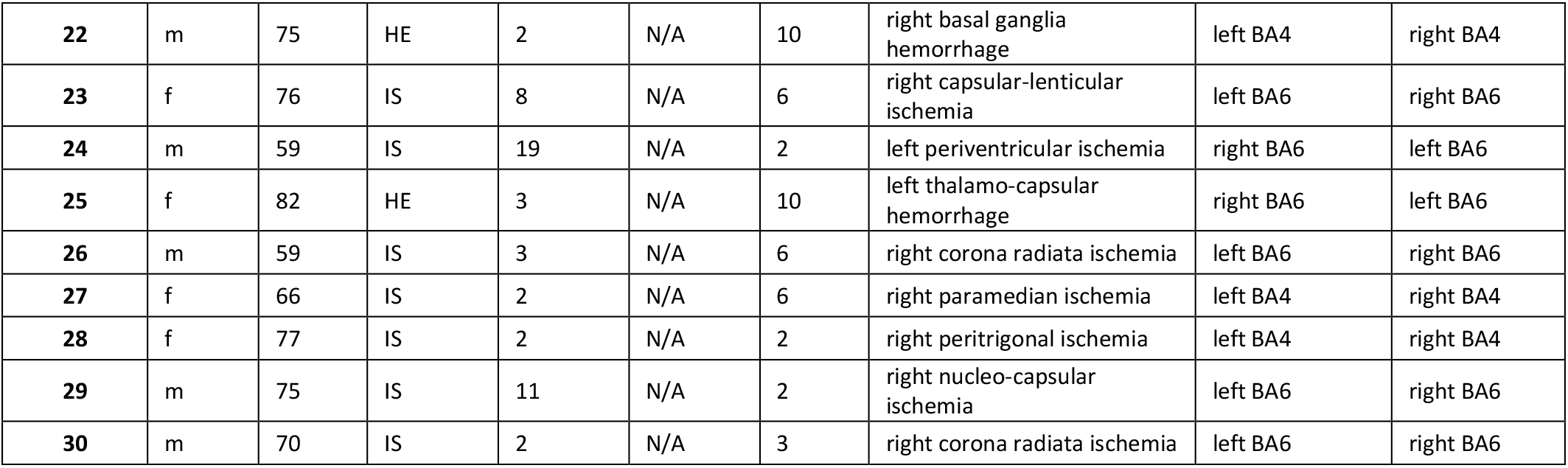
Patients demographic and clinical data. HE: Hemorrhagic. IS: Ischemic. TBI: Traumatic Brain Injury. CRS-R: Coma Recovery Scale-Revised. NIHSS: NIH Stroke Scale. BA: Brodmann Area.

Each patient underwent a single experimental session which included two TMS/EEG measurements performed with TMS targeted to intact cortical portions nearby structural lesions (perilesional stimulation site) as well as far from the lesion (contralesional stimulation site, typically located over the contralateral hemisphere as in Figure 1A, but see Materials and Methods for details regarding the specific TMS targeting, particularly regarding the severe multifocal lesions group). This experimental setting allowed for a direct within-subject control thus minimizing the variability of the dependent variables under scrutiny often present in case-control studies. Specifically, in each of the three groups, we assessed the occurrence of a TMS-evoked slow wave (<4 Hz) associated with the presence of a cortical OFF-period, i.e. a significant suppression of high frequency (>20 Hz) EEG power compared to baseline*(19, 20, 26)* confined to the perilesional stimulated site (Figure 1B,C and D). Furthermore, we aimed at assessing the effects of local cortical OFF-periods on local signal complexity as measured by the adaptation of a recently proposed index of perturbational complexity (PCI^st^;*(27)*)

**Figure 1.**
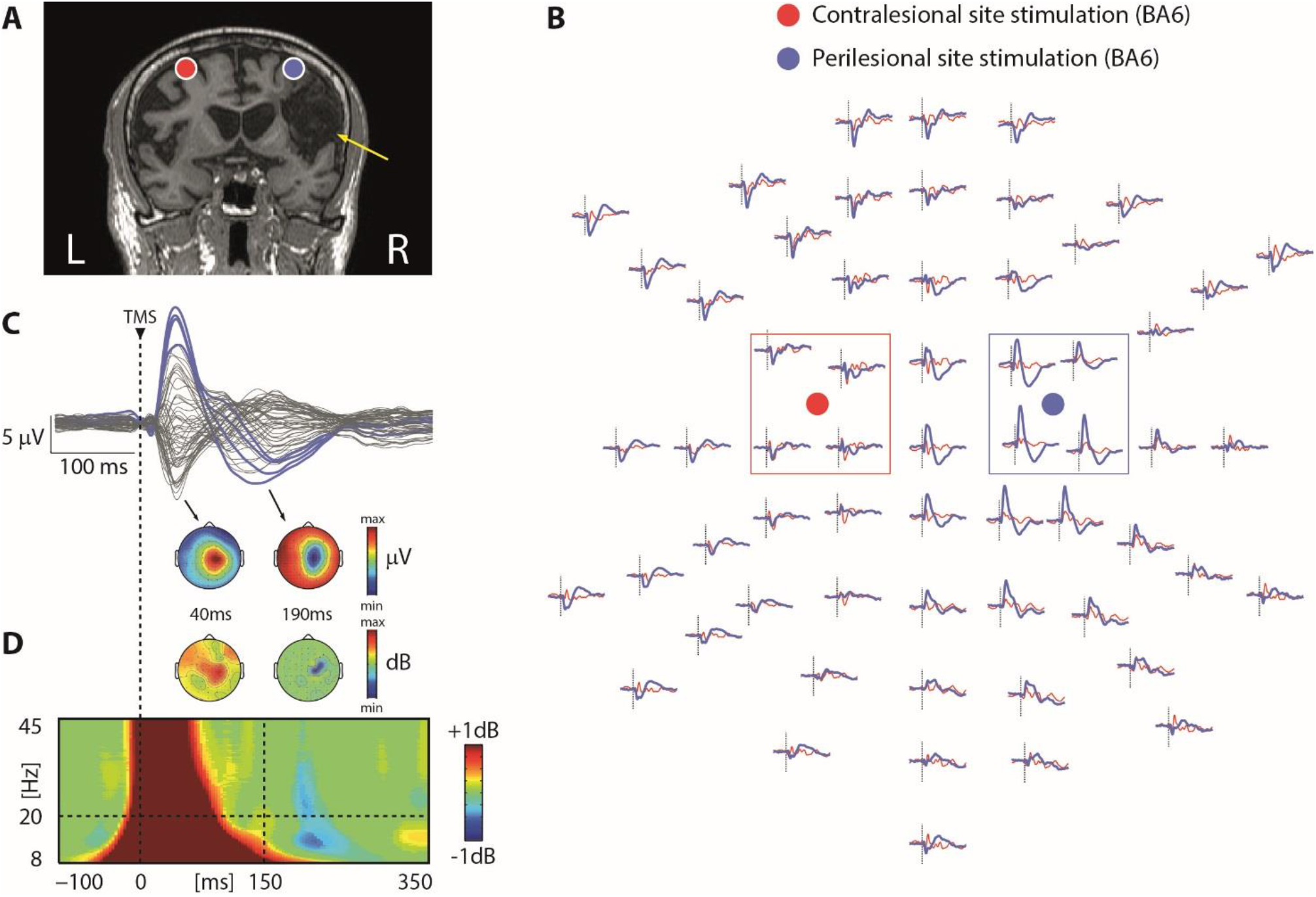
TMS reveals local, sleep-like slow waves associated with cortical OFF-periods over the affected hemisphere. Results from one representative patient (patient n.10 from Table 1) are shown for both the contralesional (red) and perilesional (blue) stimulation sites. **Panel A**. MRI and cortical targets (BA6) as estimated by the Navigated Brain Stimulation system are shown. The yellow arrow highlights lesion location. **Panel B**. Superimposition of TMS-evoked potentials of the two stimulated sites for all EEG sensors (arranged based on the channel layout is displayed. For each channel, a dashed vertical line marks the occurrence of TMS. Color-coded circles represent the position of the coil focus over the scalp. Color-coded boxes highlight the four channels closest to TMS for both stimulation sites. Note the occurrence of a local slow-wave over the right hemisphere for the perilesional stimulation site. **Panel C**. Butterfly plot of the TMS-evoked EEG potentials recorded at all 60 electrodes for the perilesional stimulation site (top). The four EEG electrode closest to TMS (indicated by the blue box in Panel B) are displayed in color. The instantaneous voltage topography of the positive and negative deflections of the slow wave is depicted below. **Panel D**. Event-related spectral perturbation (ERSP) averaged across the four EEG electrodes closest to TMS (bottom). Significance for bootstrap statistics is set at α=0.05 (absence of any significant difference from baseline spectrum is colored in green): statistically significant increases of power compared to baseline are colored in red, while blue represents significant power decreases. The dashed horizontal line indicates the 20 Hz frequency bin while the dashed vertical line at 150ms indicates the beginning of the significant high-frequency (>20 Hz) power quantification. As in Panel C, the instantaneous voltage topography of the averaged power between 20 and 45 Hz is depicted above for the same timepoints. Note the occurrence of a local significant suppression of high-frequency power limited to the perilesional stimulation site concurrent with the negative deflection of the slow wave. For Panels B C and D, a dashed vertical line at time 0 marks the occurrence of TMS.

### TMS reveals local, sleep-like cortical OFF-periods over the affected hemisphere only in patients with cortico-subcortical lesions irrespective of the etiology

In the MCA ischemia group, TMS-evoked EEG potentials (TEPs) obtained from the stimulation of the contralesional site were low-amplitude, fast-frequency recurrent scalp waves similar to those previously observed in healthy awake individuals (Figure1B and Figure 2A, red trace;*(25, 28)*). When TMS was applied using the same stimulation parameters over the perilesional site, TEPs were characterized by a local slow EEG potential (Figure 1B, C and 2A, blue traces) associated with an initial broad-band activation followed by a significant suppression of high frequency EEG power starting roughly at 150 ms after TMS (mean±SEM: 167±14 ms) over the four channels closest to the stimulation site (Figure 1B and C). Notably, this local pattern of activation, matching the criteria for an OFF-period, was found for all ten patients only over the perilesional stimulation site and never over the contralateral unaffected stimulation site. Consistently, low-frequency (<4 Hz) EEG amplitude (SWa, see Materials and Methods section) obtained over the perilesional site was significantly higher compared to the contralesional homologue site (Wilcoxon signed-rank test Z=2.4, p=0.01; Figure 2B). Also, the suppression of high frequency power (HFp, see Materials and Methods) was significantly different between the two stimulated sites (Wilcoxon signed-rank test Z=2.8, p=0.005), and only present over the perilesional stimulation site (all ten patients displayed negative values (range: −0.01/-0.6 dB, see Figure 2B). Notably, this consistent pattern of TMS-evoked response over the perilesional site was invariably found in all subjects irrespective of the presence of slow waves spontaneously occurring in their background EEG (Figure S1), and even for the two patients whose background EEG was characterized by the absence of lateralized focal anomalies (patients n.2 and n.8; Table S1).

**Figure 2.**
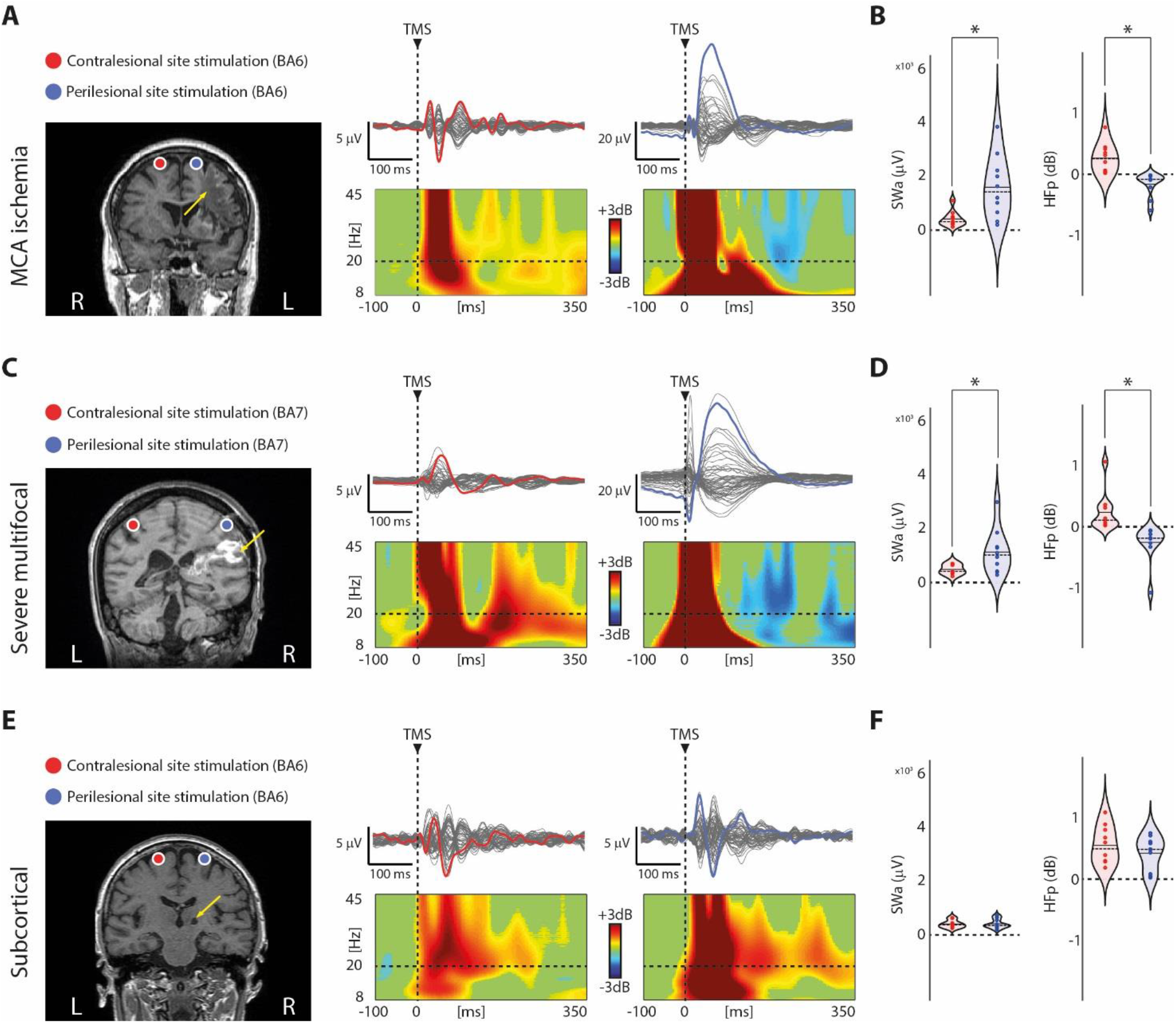
Perilesional cortical OFF-periods are present in all patients with cortico-subcortical lesions irrespective of the etiology. **Panel A, C and E**. Results from three representative patients, one for each group (patients n.4, 12 and 23 from Table 1, respectively) are shown for both contralesional (red) and perilesional (blue) stimulation sites. For each panel, MRIs and cortical targets as estimated by the Navigated Brain Stimulation system are shown (left). The yellow arrows highlight lesion location. Butterfly plots of the TMS-evoked EEG potentials recorded at all 60 electrodes (traces) are depicted (right). A dashed vertical line marks the occurrence of TMS. Event-related spectral perturbation (ERSP) is presented for the EEG electrode (colored trace) with the largest early response, selected among the four channels closest to TMS. In the ERSP plot, significance for bootstrap statistics is set at α=0.05 (absence of any significant difference form baseline spectrum is colored in green): statistically significant increases of power compared to baseline are colored in yellow/red, while blue represents significant power decreases. The dashed horizontal line indicates the 20 Hz frequency bin. **Panel B, D and F**. Violin plots and individual values of slow wave amplitude (SWa, see Materials and Methods; top) and high-frequency power (HFp, see Materials and Methods; bottom) calculated for the contralesional (red) and perilesional (blue) stimulation sites for the three groups. Violin plots display the median (dashed line) and the mean (solid line) of the kernel density. Asterisks indicate significant differences (p<0.05; Wilcoxon signed-rank test).

Similar results were obtained in the severe multifocal lesions group (Figure 2C). Also in this case, TEPs obtained from the perilesional stimulation site were characterized by a slow wave associated with an initial broad-band activation followed by a significant suppression of high frequency EEG power starting at 158±16 ms (mean±SEM) over the four channels closest to the stimulation site. Consistently (and similar to the unilateral MCA ischemia group) both SWa and HFp (Figure 2D) obtained over the perilesional site were significantly different compared to the contralesional site (Wilcoxon signed-rank test Z=2.3, p=0.01 and Z=2.8, p=0.005, respectively). Importantly, also in this group, the suppression of HFp was present for all ten patients only over the perilesional stimulation site (range: −0.06/-1.09 dB, see Figure 2D).

Overall, results in these two groups of patients confirmed that, irrespective of the etiology, TMS in the presence of cortico-subcortical lesions featured a sleep-like slow wave associated with the presence of a cortical OFF-period confined to the perilesional stimulated site. Further confirming the sleep-like nature of the observed findings, Figure S2 shows the results obtained for a representative patient (patient n.14, see Table 1) from the severe multifocal lesions group. In this patient, in addition to the two TMS/EEG measurements performed during wakefulness (Figure S2B), the same stimulations were also performed while the patient was asleep (Figure S2B) for the entire duration of the recordings (as assessed by the presence of prolonged eye closure and typical NREM sleep graphoelements -i.e. sleep spindles and slow waves-in the spontaneous EEG). Results show a striking similarity between the TMS-evoked response obtained over the perilesional stimulation site during wakefulness and those obtained over both stimulation site during sleep, particularly with respect to the presence of significant high-frequency power suppression concurrent with the negative deflection of the slow wave (the hallmark of cortical OFF-periods during sleep).

At odds with the above, TEPs obtained in the subcortical lesions group (Figure 2E) were never associated with the presence of a clear-cut slow wave or high frequency EEG power suppression over neither the perilesional nor the contralesional stimulation site (Wilcoxon signed-rank test Z= 0.7, p=0.44 for SWa and Z=0.8, p=0.38 for HFp, see Figure 2E and F). Interestingly, for both SWa and HFp, the values obtained in this group of patients over both sites were similar to those obtained over the contralesional stimulation site in both MCA ischemia and severe multifocal lesions groups (Figure 2B, D and F). To test this, we performed a mixed-model ANOVA with GROUP as categorical predictor and SITE as within-subject factor. For both SWa and HFp, values obtained in the subcortical lesions group over both perilesional and contralesional sites were not statistically different from those obtained over the contralesional site of both MCA ischemia and severe multifocal lesions, as revealed by post-hoc comparisons (all p-values >0.2, Bonferroni corrected). To further explore this, in one patient affected by multiple lacunar periventricular ischemic white matter lesions (patient n.24, see Table 1) we performed a more extensive mapping including two additional stimulations over BA 7 of both the affected and the unaffected hemispheres (Figure S3). Results confirmed the absence of TMS-evoked slow waves and OFF-periods irrespective of the stimulated hemisphere and cortical area.

### The presence of local, sleep-like cortical OFF-periods affects local signal complexity

We then assessed the effects of the occurrence of perilesional TMS-evoked slow waves and OFF-periods on local signal complexity. To this aim, for each patient and stimulated site, we quantified the temporal complexity of the principal components of the signals calculated over the four channels closest to TMS (Figure 3A, C and E). The complexity of the local TMS-evoked responses was found significantly reduced over the perilesional compared to the contralesional stimulation site for the MCA ischemia group (Wilcoxon signed-rank test Z=2.4, p=0.01; Figure 3B). A similar, albeit not statistically significant result was found for the severe multifocal lesions group where differences in local signal complexity between stimulation sites were only showing a statistical trend (Wilcoxon signed-rank test Z=1.5, p=0.1; Figure 3D). The same analysis applied on the subcortical lesions group showed no differences in local signal complexity between stimulation sites (Wilcoxon signed-rank test Z=1.2, p=0.2; Figure 3F). Altogether, these findings suggest that, when present, local sleep-like cortical OFF-periods affect local signal complexity.

**Figure 3.**
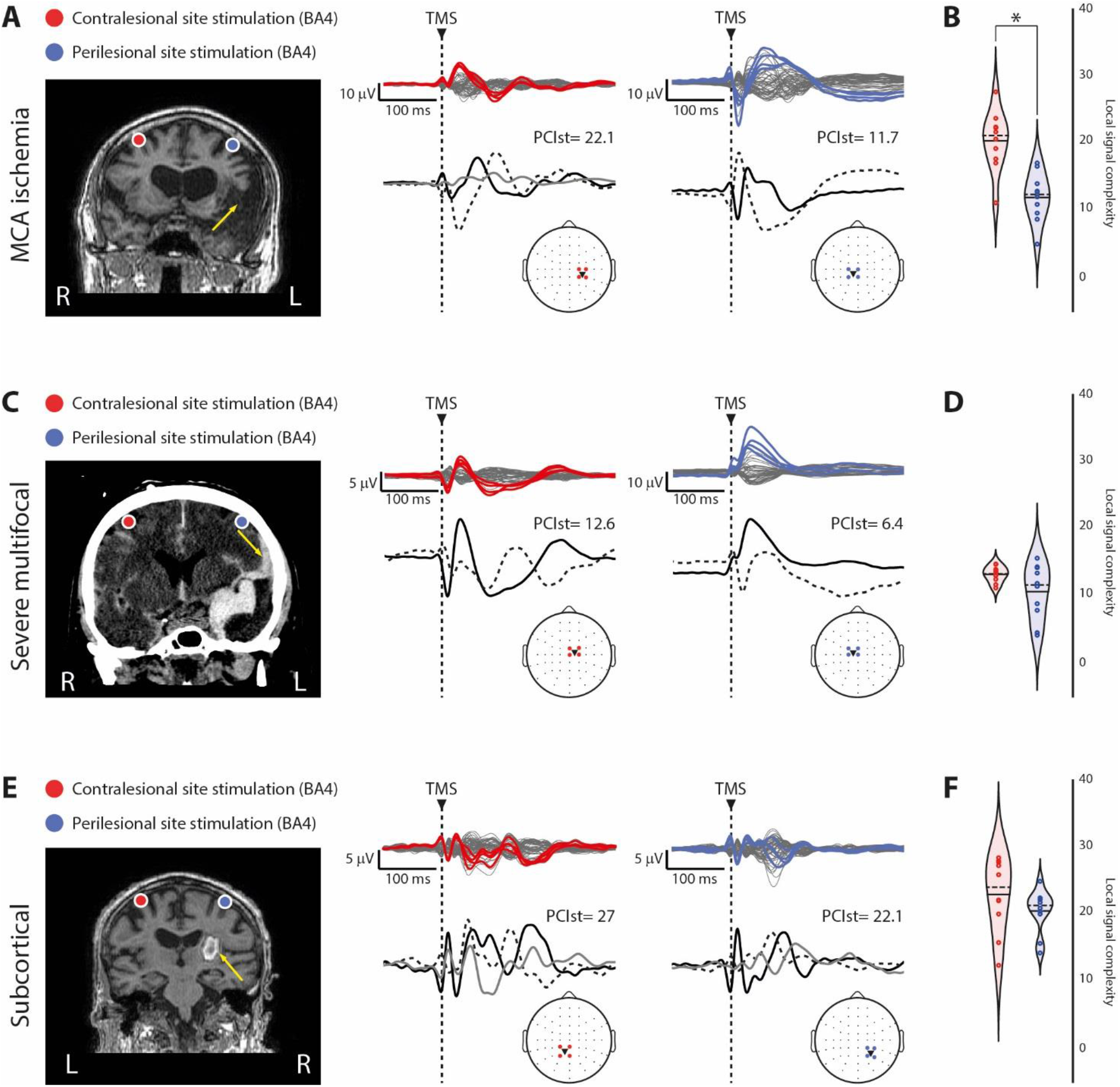
The presence of local, sleep-like cortical OFF-periods affects local signal complexity. **Panel A, C and E**. For each panel, brain imaging (MRI for Panel A and E, CAT scan for Panel C) and cortical targets as estimated by the Navigated Brain Stimulation system are shown for three representative patients, one for each group of patients (patients n.1, 13 and 22 from Table 1, respectively) are shown for both the contralesional (red) and perilesional (blue) stimulation sites. The yellow arrows highlight lesion location. Butterfly plots of the TMS-evoked EEG potentials recorded at all 60 electrodes (traces) are depicted (top). For each stimulated site, the four EEG electrode closest to TMS are displayed in color and topographically projected on the template channel layout (bottom). The correspondent local signal complexity values (calculated by applying PCI^st^ restricted to the four channels, see Materials and Methods) and the time-course of the principal components based on singular value decomposition are depicted (bottom). A dashed vertical line marks the occurrence of TMS. **Panel B, D and F**. Violin plots and individual values of local signal complexity calculated for the contralesional (red) and perilesional (blue) stimulation sites for the three groups. Violin plots display the median (dashed line) and the mean (solid line) of the kernel density. Asterisks indicate significant differences (p<0.05; Wilcoxon signed-rank test).

Of note, the somehow smaller difference observed in the group of patients affected by severe multifocal lesions may be explained by the presence of brain lesions often involving both hemispheres (see Table 1), partially affecting also the local signal complexity of the contralesional stimulation site. This was confirmed by the marginal effect found for the mixed-model ANOVA with GROUP as categorical predictor and SITE as within-subject factor (GROUP*SITE interaction effect: F(2, 27)=3.3, p=0.04985). In particular, regarding the perilesional stimulation site, local signal complexity for the severe multifocal lesions group was found similar to that observed for the MCA ischemia group (p>0.05, Bonferroni corrected; Figure 3B and D) and significantly lower than that observed in the subcortical lesions group (p=0.000001, Bonferroni corrected; Figure 3D and F). Regarding the contralesional stimulation site, local signal complexity was found reduced for the severe multifocal lesions group compared to the other two groups (all p-values <0.0002, Bonferroni corrected; Figure 3B, D and F).

Further we assessed the impact of focal brain injuries on the global spatiotemporal dynamics of TEPs, which reflect the overall capacity of thalamocortical circuits to engage in complex patterns of causal interactions, typically present in conscious awake individuals*(27, 29)*. Thus, for each patient we estimated the maximum PCI^st^ calculated on all the sixty channels following the stimulation of the contralesional site. Results at the group level were in line with previous observation*(30)* in conscious brain-injured individuals and showed that, for all three groups, average maximum PCI^st^ values (mean ± SE: 41.1 ± 2.4, 28.5 ± 3.1 and 42.5 ± 2, for the MCA ischemia, the severe multifocal and the subcortical lesion groups, respectively; Table S2) were above the empirical threshold (PCI^st^ threshold value: 23.02;*(27)*) for consciousness obtained from a benchmark population (382 TMS/hd-EEG sessions performed on 108 healthy subjects). However, a ONE-way ANOVA (GROUP effect F(2, 27)=9.3, p=0.00082) showed a significantly lower maximum PCI^st^ value for the severe multifocal lesions group compared to the other two groups (all p-values <0.004, Bonferroni corrected), confirming the role of multifocal brain lesions in partially affecting overall signal complexity (PCI^st^ was found lower than the benchmark empirical threshold in patients n.17 and 18, gray text in Table S2).

## Discussion

This study provides a novel insight about the nature of the electrophysiological consequences of brain injury. Reviving the old notion of EEG slowing in perilesional areas, here we draw an explicit connection between the electrophysiology of local sleep, thoroughly described in the sleep literature, and the pathophysiology of focal brain injury and stroke.

In doing this, the present work also reinforces and generalizes the idea that TMS/EEG represents an interesting venue for the identification of relevant markers of recovery after stroke*(31–35)*. Indeed, the current finding that local sleep-like slow waves and OFF-periods characterize the cortical tissue surrounding the injury of different brain areas expands the potential of TMS/EEG beyond the motor domain (Figure 1, Figure 2).

The hallmark of physiological sleep is the occurrence of slow waves and OFF-periods, often referred to in the sleep literature as cortical bistability*(7, 10)*. OFF-periods are caused by the enhancement of adaptation (or activity-dependent) K+ currents, brought about by decreased levels of neuromodulation from brainstem activating systems*(36–39)* and/or by increased inhibition*(40–42)*. Due to these mechanisms, cortical neurons tend to plunge into a silent, hyperpolarized state, lasting few hundred milliseconds, after an initial activation*(10)*. The occurrence of synchronous membrane hyperpolarization and synaptic silence in cortical neurons is reflected in extracellular slow waves associated with transient suppressions of high-frequency (>20 Hz) activity that may be detectable both in the local field potential*(19, 20)* and in the EEG*(26, 43)* spontaneous activity. However, due to its activity-dependent nature, cortical bistability and the associated OFF-periods can be better revealed using a perturbational approach, whereby the impulse-response properties of cortical neurons are challenged by means of direct activations*(22, 23)*. Consistently, TMS-evoked EEG events similar to those reported here were previously found in sleeping*(25, 44, 45)* and anesthetized*(46, 47)* healthy controls and never in healthy awake subjects*(25, 28)*.

A key finding of the present work is the demonstration of a pathological form of local cortical bistability sharing similar features to that characterizing the sleep state (see Figure S2) but occurring (i) during wakefulness, and (ii) in circumscribed intact portions of the cortex adjacent to focal brain injuries.

Interestingly, in the last few years, science has produced compelling evidence supporting the idea that sleep is under local regulation*(48)*. Local sleep is a complex physiological phenomenon occurring within anatomically discrete brain locations. Experiments in isolated cortical slabs*(49)*, as well as in slice preparations*(7)* and cell cultures*(50, 51)*, confirmed that sleep— in the form of the occurrence of slow waves, the key electrophysiological graphoelement characterizing the sleep state—is essentially an intrinsic property of cortical cells ensembles*(52)*. In parallel, a growing body of evidence, expanded the notion of local sleep partially redefining the classical definition of wake and sleep as separate, discrete states. A number of studies in both rodents and humans demonstrated the occurrence of local sleep-like graphoelements intruding behavioral wakefulness over circumscribed cortical regions*(14–18)*. A number of factors may account for such occurrence and include a regional-specific sensitivity to the activating signals from the brainstem and hypothalamic afferents*(53, 54)*, the diverse influence of the homeostatic process*(55)* over different brain structures and/or variable degrees of synaptic strength differentially impinging on local circuits in a use-dependent manner*(56)*.

Particularly relevant in the context of the present manuscript is that local sleep may also be promoted by cytokines such as interleukin-1 (IL-1) and tumor necrosis factor α (TNFα)*(57)* which enhance hyperpolarizing inward Cl-currents*(58)*, and outward K+ currents*(59)*, respectively. Notably, local cortical unilateral microinjections of both these cytokines in rats result in an acute lateralized increase in the number of EEG slow waves during subsequent sleep*(60, 61)*. Similarly, the local immune response taking place over perilesional areas may be responsible for the occurrence of the lateralized EEG slow waves typically observed in the acute post-stroke phase*(6)*.

In parallel, the reduction of the ATP/ADP ratio following ischemia, activates K-ATP sensitive channels (a class of ionic channels conducting inward-rectifier K+ currents) ubiquitously expressed in neuronal*(62)* as well as in glial cells*(63)*. Their activation causes membrane hyperpolarization and suppression of activity in local neuronal populations surrounding the lesion*(64)*. This phenomenon, although initially neuroprotective, may have chronic functional consequences on the cortical tissue surviving the insult depending on the length of the hypoxic challenge. Specifically, after prolonged hypoxia, the cell membrane potential may not follow the typical speedy recovery and stabilization*(64)*, thus resulting in a prolonged hyperpolarization in local cortical circuits surrounding the lesion. In this case, the surviving neuronal populations would be chronically characterized by sleep-like excitability profiles similar to those observed in the present work.

Besides the local effects of inflammatory and metabolic changes, pathological alterations of the excitation/inhibition balance similar to those occurring during physiological sleep*(10, 65)* may play a major role after brain injury. Specifically, in a mouse model of stroke*(12)*, tonic neuronal inhibition was found increased in the peri-infarct zone. Notably, dampening tonic inhibition by means of a benzodiazepine inverse agonist produced an early and sustained recovery of motor function. Consistently, recovery of language and motor function after stroke can be blocked by a chronic excessive inhibitory activity over the perilesional areas*(31)*. On the other hand, lesions may induce local cortical bistability by engendering a state of cortico-cortical disfacilitation, that is by reducing recurrent excitation*(66)* from the lost tissue. Since cortical neurons receive most of their inputs from nearby cells, they tend to become hyperpolarized when their neighbors drastically reduce their spiking activity (e.g. during sleep*(67)*). Cortical lesions of various etiologies (including traumatic) represent an extreme case of such, resulting in an intracortical fiber-mediated hyperpolarization and depression of function in intact cortical areas neighboring the injured site, i.e. *diaschisis associativa (1)*.

Cortical bistability has been also experimentally induced in the animal model by means of cortical deafferentation*(66, 68)*. As an example, severing the white matter with a cortical undercut results in slow waves and in a continuous alternation between ON and OFF-periods in the partially deafferented gyrus, even when the animal, and the rest of the brain, is awake*(11)*. To the extreme, cutting a slice of cortex and bathing it in the appropriate in vitro solution results in a typical pattern characterized by the alternation between ON- and OFF-periods*(9)*. Deafferentation may thus induce cortical bistability due to the interruption of a critical amount of fibers from the ascending activating systems and may occur in patients following both large vascular (e.g. MCA ischemia) and traumatic (diffuse axonal injury) white matter lesions.

Irrespective of the specific mechanism involved, direct cortical perturbations with TMS could best reveal the presence of local OFF-periods regardless of the background EEG pattern, of the presence of focal anomalies (e.g. absent in patient n.2 and n.8) and, most importantly, of the prevalence of spontaneously lateralized slow waves (found, with variable incidence, in five out of ten MCA ischemia patients). This evidence clearly highlights the potential of TMS/EEG as an informative investigational tool for studying the electrophysiological consequences of focal brain lesions above and beyond spontaneous EEG*(69)* (Table S1 and Figure S1). Indeed, a direct cortical hit with TMS may best reveal the presence of covert forms of bistability by (i) massively recruiting local inhibitory circuits if the excitation/inhibition balance is biased towards the latter, as well as (ii) trigger activity-dependent K+ currents if K+ channels are de-inactivated.

In this context, the cortical involvement characterizing the lesions of both MCA ischemia and severe multifocal patients, is consistent with all the mechanisms outlined above. Also, the absence of EEG signs of cortical bistability in the group of patients affected by unilateral subcortical lesions further supports these mechanistic interpretations. Indeed, the occlusion/rupture of small penetrating arteries characterizing lacunar ischemic as well as hemorrhagic subcortical lesions share distinct features with respect to the occlusion of large vessels, i.e. MCA ischemia, and to multifocal lesions of various etiologies. Specifically, purely subcortical lesions tend to (i) be smaller in volume and (ii) involve efferent pathways and/or structures. As such, lesion extent and location may not be optimal for chronically altering the excitability profiles of connected cortical areas through the above-mentioned mechanisms. This interpretation is in line with classic reports of an absence of lateralized EEG slow waves following lacunar infarction*(70)* and is further confirmed by TMS/EEG work in patients with unilateral subcortical lesions of the internal capsule, thalamus, and basal ganglia. This work reported the occurrence of EEG responses characterized by a sequence of positive and negative polarity deflections starting a few milliseconds post-stimulation with similar morphology compared to healthy controls*(35)*. The same study also reported a reduced amplitude of fast frequency oscillatory components in the group of patients. Similarly, in our subcortical lesion group the frequency of the oscillatory components (i.e., the natural frequency*(28)*) was found reduced for both stimulated sites compared to that of healthy subjects reported in previous work*(71)* for the same stimulation sites (Table S3).

Ultimately, the presence of OFF-periods in areas that are not directly affected by the lesion may have profound behavioral consequences. In this context, increasing evidence suggest that additional symptoms following brain injury actually result from the local functional disruption in intact brain areas anatomically and functionally connected to those affected by the insult*(1–5)*. In an extreme case, the ubiquitous presence of pathological OFF-periods in response to TMS across cortical islands spared by anatomical lesions in awake VS/UWS patients was shown to prevent the build-up of global complexity, resulting in behavioral unresponsiveness*(25, 29, 30)*. Here, we report for the first time the presence of three different local patterns of cortical reactivity within the same patient (Figure S4). Specifically, the response was absent when directly stimulating the lesion, sleep-like over the perilesional areas, and typical of wakefulness over contralateral areas. Interestingly, the occurrence of local OFF-periods over perilesional areas was associated with the disruption of local signal complexity (Figure 3) - global complexity being in the typical range of conscious awake individuals (Table S2) - which may impair specific behavioral domains. Establishing a systematic link between the presence of local OFF-periods in the perilesional area and their behavioral consequences would ultimately require longitudinal measurements paralleled by an appropriate domain-specific clinical assessment, which represents a limitation of the present study. However, previous studies performed in sleep-deprived rodents*(15)* and humans*(18)* have shown that, even in the absence of lesions, the intrusion of local sleep-like electrophysiological features in the awake brain are associated with selective motor impairment or cognitive lapses depending on the specific cortical regions involved.

Overall, these findings directly connect the field of brain injury to the rich wealth of knowledge that we have accumulated about the cellular and network mechanisms of slow waves. To the extent that local sleep-like neuronal bistability represents the key electrophysiological alteration of perilesional cortical areas, OFF-periods may represent a valid read-out of the state of discrete cortical circuits surviving the insult, as well as a potential target for the development of novel therapeutic interventions aimed at restoring a more balanced pattern of activation. An intriguing possibility is that intact cortical areas that are active and reactive may be stuck in a chronic, yet reversible, sleep-like regime. If confirmed by future longitudinal studies, reducing perilesional cortical bistability may then, in turn, help fostering the recovery of function in brain-injured individuals.

## Materials and Methods

### Patients

Ten patients (4 F; age [y ± SEM]: 68 ± 4) affected by cortico-subcortical lesions due to an ischemic occlusion of the MCA as well as ten patients (4 F; age [y ± SEM]: 54 ± 6.1) affected by severe multifocal cortico-subcortical lesions of various (ischemic, haemorrhagic or traumatic) aetiologies and ten patients (6 F; age y ± SEM: 72 ± 2.6) affected by unilateral purely subcortical lesions (either lacunar ischemic stroke or typical haemorrhage) in a sub-acute to chronic stage (>1 months; mean duration in months ± SEM: 12.9 ± 3.9; 7.6 ± 2.5; 19.5 ± 8.2, respectively) were included in the study. For the MCA ischemia and subcortical lesion groups, clinical evaluation included the NIH Stroke Scale (NIHSS;*(72)*) (average [min/max] NIHSS score: 8 [4/17] and 5.5 [2/10], respectively; Wilcoxon rank sum test Z=2.1, p=0.03). The Coma Recovery Scale-Revised (CRS-R;*(73)*) was applied for the clinical evaluation of the severe multifocal lesions group which included patients in a Minimally Conscious State (MCS; n=6) as well as patients who emerged from the MCS and recovered functional communication (EMCS; n=4).

Anatomical lesions were assessed by means of T1-weighted Magnetic Resonance Imaging (MRI) or Computerized Axial Tomography (CAT) scans acquired before the experimental sessions. For all patients, exclusion criteria were: positive remote or familiar history for epilepsy and/or seizures, positive remote or familiar history for convulsive events, presence of metallic cranial implants, presence of TMS incompatible equipment (pace-maker, drugs dispenser, cochlear implants, intracranial stimulator), pregnancy, personal history for alcohol or drug abuse and/or for psychiatric conditions.

The experimental protocols were approved by the local ethical committees of the following Institutions: Istituto di Ricovero e Cura a Carattere Scientifico Fondazione Don Gnocchi Onlus, Fondazione Europea per la Ricerca Biomedica Onlus in Milan, Italy, and the Faculty of Medicine of the University of Liege in Liege, Belgium. A written informed consent was obtained from all the patients or their legal surrogates (as in the case of MCS patients pertaining to the severe multifocal lesions group).

### Experimental procedures

During the entire duration of the experiment, patients were awake and with their eyes open. If signs of drowsiness appeared, recordings were momentarily interrupted and subjects were stimulated using the CRS-R arousal facilitation protocols*(73)*.

The presence of cortical lesions based on the individual anatomical T1-weighted MRI or CAT scans guided the selection of TMS targets*(25, 74)*. Images were acquired for all the patients included in the study within 1 week prior to the TMS/EEG assessment. Specifically, depending on the location and spatial extent of lesions in each individual patient, we included in the present study the analysis of two TMS/EEG measurements (see below for details regarding the specific TMS targeting). The precision and reproducibility of the TMS pulses with respect to the selected targets was guaranteed by means of a Navigated Brain Stimulation (NBS) system (Nexstim Ltd., Finland).

For each of the three patient groups, the stimulation sites were operationalized as follows. In the case of ischemic unilateral MCA stroke patients, the perilesional stimulation site corresponded to the intact portion of the BA affected by the lesion (either BA4, BA6 or BA7) while the contralesional stimulation site to the homologue contralateral cortical area. In the case of patients affected by severe multifocal lesions we followed the same criteria applied for the cortical lesion patient group and we targeted TMS over the intact portion of the same BA affected by the lesion (either BA4, BA6 or BA7; perilesional stimulation site) as well as over the homologue contralateral cortical area (contralesional stimulation site), unless also directly affected by a lesion itself or inaccessible to TMS due to the presence of intracerebral drainage/shunt. In this case, the contralesional stimulation site was chosen as a BA spared from lesions/shunt either over the same (patients n.16 and 18) or the contralateral (patients n.11 and 19) hemisphere. Finally, for unilateral subcortical stroke patients, the perilesional stimulation site consisted in a frontal (either BA4 or BA6) target over the affected hemisphere, while the contralesional stimulation site to the homologue contralateral cortical area. Of note, for all patients, when targeting TMS to BA4 we explicitly avoided stimulating the hand area in order to avoid contamination by the sensory re-entry of proprioceptive feedback associated with target muscle activation, which have been previously shown to affect the spectral features of TEPs*(75)*. A detailed description regarding the lesions as well as the contralesional and perilesional stimulation sites for each individual patient are shown in Table 1.

Stimulation pulses were delivered with a Focal Bipulse figure-of-eight coil (mean/outer winding diameter ~50/70 mm, biphasic pulse shape, pulse length ~280 μs, focal area of the stimulation 0.68 cm^2^) driven by a Mobile Stimulator Unit (eXimia TMS Stimulator, Nexstim Ltd., Finland). For all the TMS/EEG measurements, the location of the maximum electric field induced by TMS on the cortical surface (hotspot) was always kept on the convexity of the targeted cortical gyrus with the induced current perpendicular to its main axis. Each cortical target was stimulated with an estimated electric field, orthogonal to the gyral crown, of about 120V/m corresponding to a percentage of the maximal stimulator output (% MSO) intensity comparable between contralesional and perilesional stimulation sites for each patient group (mean ± SEM: 65.3 ± 1.97 vs 67.1 ± 2.61 for the MCA ischemia group, 62.6 ± 3.7 vs 67.8 ± 2.58 for the severe multifocal lesions group and 59.2 ± 1.73 vs 60.9 ± 2.24 for the subcortical lesion group; for all comparisons p>0.05, paired t-test). In each TMS/EEG measurement, at least 200 stimulation pulses were delivered with an interstimulus interval randomly jittering between 2000 and 2300 ms (0.4–0.5 Hz). EEG data were recorded using a TMS-compatible 60-channel amplifier (Nexstim Ltd, Finland), which gates the magnetic pulse artefact and provides artifact-free data from 8 ms after stimulation*(76)*. Raw recordings were online referenced to an additional forehead electrode, filtered between 0.1-350 Hz and sampled at 1450 Hz. Two additional sensors were applied to record the electroculogram (EOG). As previously recommended*(77)*, during all TMS/EEG recordings a masking sound was played via earphones and a thin layer of foam was placed between coil and scalp for abolishing the auditory potentials evoked by TMS coil’s discharge. Data analysis was performed using Matlab R2012a (The MathWorks Inc.).

### Resting-state EEG data recording and analysis

Preceding TMS/EEG measurements and using the same EEG recording apparatus, a wake resting state hd-EEG recording (up to 10 minutes with eyes open) was performed for the MCA ischemia patients (n=10) to assess the presence of the lateralized slowing in the theta/delta frequency range typical of unilateral brain injuries characterized by cortical infarction*(70)*.

Spontaneous EEG data were offline filtered (0.5–40 Hz) with a 3rd order IIR Butterworth digital filter with an attenuation of −3dB at 0.5 and 40 (using the filtfilt function in the MATLAB signal processing toolbox). Continuous data were then split into contiguous 2-second segments. Artifactual segments were excluded from the analysis based on visual inspection (max/min retained EEG segments: 284/69). Bad channels were rejected based on visual inspection (≤ 10% of channels per recording). Rejected channels were then interpolated using spherical splines. The signal for each channel was then re-referenced to the average of all channels. After reducing the number of independent components to the number of good, non-interpolated channels by performing Singular Value Decomposition, ICA was used to remove ocular, muscle, and cardiac pulse artifacts using EEGLAB routines. For each EEG derivation, power spectral density (PSD) estimates were computed using the Welch’s method with 2-s Hanning windows and 50% overlap (applying the *pwelch* function in the MATLAB signal processing toolbox). For each patient, average PSD was calculated over the same four channels used for TMS/EEG analysis (contralesional, perilesional): average PSD across segments was computed for each frequency bin and then further averaged across bins pertaining to the classical frequency ranges: delta (0.5-4.5Hz), theta (4.5-8 Hz), alpha (8-12 Hz), beta (12-30 Hz) and gamma (30-40 Hz) (Figure S1A). In parallel to the PSD analysis, filtered continuous EEG data were re-referenced according to a longitudinal bipolar montage based on the 10-20 system*(78)* including the following EEG derivations: ‘Fp1_F5’, ‘F7_T3’, ‘T3_T5’, ‘T5_O1’ ‘Fp1_F3’, ‘F3_C3’, ‘C3_P3’, ‘P3_O1’, ‘Fz_Cz’, ‘Cz_Pz’, ‘Fp2_F6’, ‘F6_T4’, ‘T4_T6’, ‘T6_O2’, ‘Fp2_F4’, ‘F4_C4’, ‘C4_P4’, and ‘P4_O2’. These data were then visually inspected by a clinical neurophysiologist to assess the presence of EEG anomalies (Table S1 and Figure S1B).

### TMS/EEG data recording and analysis

TMS/EEG recordings were visually inspected to reject trials and channels containing noise or muscle activity as in previous works*(25, 30, 74)*. Then, EEG data were band-pass filtered (1-45 Hz, Butterworth, 3rd order), down-sampled to 725 Hz and segmented in a time window of ± 600 ms around the stimulus. Bad channels (≤ 10% of channels per recording) were interpolated using spherical splines as implemented in EEGLAB*(79)*. Then, a comparable number of good trials between contralesional and perilesional stimulation sites for each patient group (mean ± SEM: 184 ± 16 vs 184 ± 14 for the MCA ischemia group, 150 ± 12 vs 137 ± 13 for the severe multifocal lesion group and 184 ± 20 vs 196 ± 23 for the subcortical lesion group; for all comparisons p>0.05, paired t-test) were re-referenced to the average reference and baseline corrected. Finally, after reducing the number of independent components to the number of good, non-interpolated channels by performing Singular Value Decomposition, Independent Component Analysis (ICA) was applied in order to remove residual eye blinks/movements and spontaneous scalp muscle activations.

In order to detect the presence of local sleep-like activity in the perilesional areas of awake, conscious brain injured patients, we followed the same methodological rationale as in*(22)* and*(25)*. Specifically, for each group of patients (n=10 each), we compared the TMS/EEG measurements performed over the contralesional and the perilesional stimulation sites and we aimed at quantifying 1) the presence of TMS-evoked slow waves over the perilesional area as well as 2) the correspondent occurrence of a cortical OFF-period. Operationally, these variables can be quantified respectively as 1) the amplitude of low-frequency EEG components (< 4 Hz), and 2) the modulation of post-stimulus high-frequency EEG power (> 20 Hz)*(19, 20, 26)*. We refer the reader to*(22, 25)* for detailed methodological description. In brief, to assess the presence of TMS-evoked slow waves, single trials were low-pass filtered below 4 Hz (second-order Chebyshev filter), re-referenced to the mathematically-linked mastoids, averaged and rectified. For each channel, the Slow Wave amplitude (SWa) was computed as the cumulated amplitude of the rectified signal within the 8-350 ms time window. On the other hand, to assess the presence of the correspondent cortical OFF-period, we applied the event related spectral perturbation (ERSP) procedure implemented in EEGLAB*(79)*. Specifically, single trials were time-frequency decomposed between 8 and 45 Hz using Wavelet transform (Morlet, 3.5 cycles; as in*(28)*) and then normalized with the full-epoch length (here ranging from −350 to 350 ms) single-trial correction*(80)*. The resulting ERSPs were averaged across trials and baseline corrected (from −350 to −100 ms). Furthermore, power values that were not significantly different from baseline (bootstrap statistics, α=0.05, number of permutations = 500) were set to zero. High-frequency EEG power (HFp) was obtained as the integral of the significant high-frequency (>20 Hz) power between 150 and 350 ms. For each TMS/EEG measurement, SWa and HFp were computed at the single-channel level and then averaged over the four channels closest to the stimulation site*(25, 81)* for group analysis.

For each patient, we then first estimated the maximum global spatiotemporal dynamics of the TMS-evoked response by computing the Perturbational Complexity Index based on the quantification the temporal complexity of the principal components of the EEG response to TMS (PCI^st^) as described in*(27)* (Table S2). In addition, we applied a generalization of the original method and, for each stimulation session, we calculated PCI^st^ restricted to the four channels closest to the stimulation site. In this way, we were able to assess the impact of perilesional sleep-like OFF-periods on local signal complexity.

In general, PCI^st^ is an index that combines dimensionality reduction and a novel metric of recurrence quantification analysis (RQA) to empirically quantify perturbational complexity as the overall number of non-redundant state transitions caused by the perturbation. Briefly, principal component analysis (PCA) of the response is performed in order to obtain the spatial modes of the signal and a method derived from RQA is then applied to quantify the number of “state transitions” present in each component. For both the global and local signal complexity estimates, PCI^st^ was computed using the source-code available at github.com/renzocom/PCIst using the same parameters for component selection (max_var=99%; min_snr=1.1) and state transition quantification (k=1.2) as in*(27)*.

### Statistical analyses

For SWa, HFp, local signal complexity, as well as for resting-state EEG PSD, within-group comparisons between stimulation sites (perilesional, contralesional) were performed by means of the non-parametric Wilcoxon signed-rank test (α=0.05; n=10). When testing between-group differences, either mixed-model ANOVAs were performed with GROUP as categorical predictor and SITE as within-subject factor (α=0.05; n=30) with post hoc two-tailed t tests (α=0.05, Bonferroni correction), or Wilcoxon rank-sum test (α=0.05; n=20) were used.

## Acknowledgments

The authors thank Dr. Angela Comanducci, Dr. Andrea Pigorini, Dr. Ezequiel Mikulan, and Mr. Simone Russo for their help and comments on the manuscript draft. This research was supported by the European Union’s Horizon 2020 Framework Program for Research and Innovation under the Specific Grant Agreement No. 720270 (Human Brain Project SGA1) (to M.M., M.R., and S.L.) and No. 785907 (Human Brain Project SGA2) (to M.M., M.R., and S.L.) and by the EU grant H2020, FETOPEN 2014-2015-RIA no. 686764 “Luminous” (to M.M. and S.L.). The study has been also funded by the grant “Sinergia” CRSII3_160803/1 of the Swiss National Science Foundation (to M.M.), by the James S. McDonnell Foundation Scholar Award 2013 (to M.M.), by the Tiny Blue Dot Foundation (to M.M.), by the Belgian National Funds for Scientific Research (F.R.S-FNRS; to S.L. and O.G.), by the Fondazione Europea di Ricerca Biomedica (to S.L. and O.G.), by the BIAL Foundation (to S.L. and O.G.), by AstraZeneca (to S.L. and O.G.) and by the Foundation Roi Baudouin (to S.L. and O.G.).

## Notes

#### Summary of Updates

Now including Acknowledgments

