## Supplementary material for "Local sleep-like cortical reactivity in the awake brain after focal injury": FigureS1; FigureS2; FigureS3; FigureS4; TableS1; TableS2; Table S3

**Fig. S1.** Resting-state EEG recordings in the MCA ischemia group.

**Fig. S2.** The TMS-evoked response over the perilesional stimulation site during wakefulness is similar to the typical TMS response ubiquitously observed during NREM sleep.

**Fig. S3.** The absence of TMS-evoked slow waves and OFF-periods irrespective of the stimulated hemisphere and cortical area in the group of patients affected by unilateral lacunar ischemic or haemorrhagic subcortical lesions.

**Fig. S4.** TMS reveals three distinct cortical reactivity profiles within the same patient.

**Table S1.** Spontaneous EEG clinical assessment of MCA ischemia patients.

**Table S2.** Maximum global spatiotemporal dynamics of the TMS-evoked responses assessed via the Perturbational Complexity Index ($PCI^{st}$ (27)).

**Table S3.** Assessment of the frequency of the main oscillatory components of the TMS-evoked response (natural frequency (28)) in the subcortical lesion group.

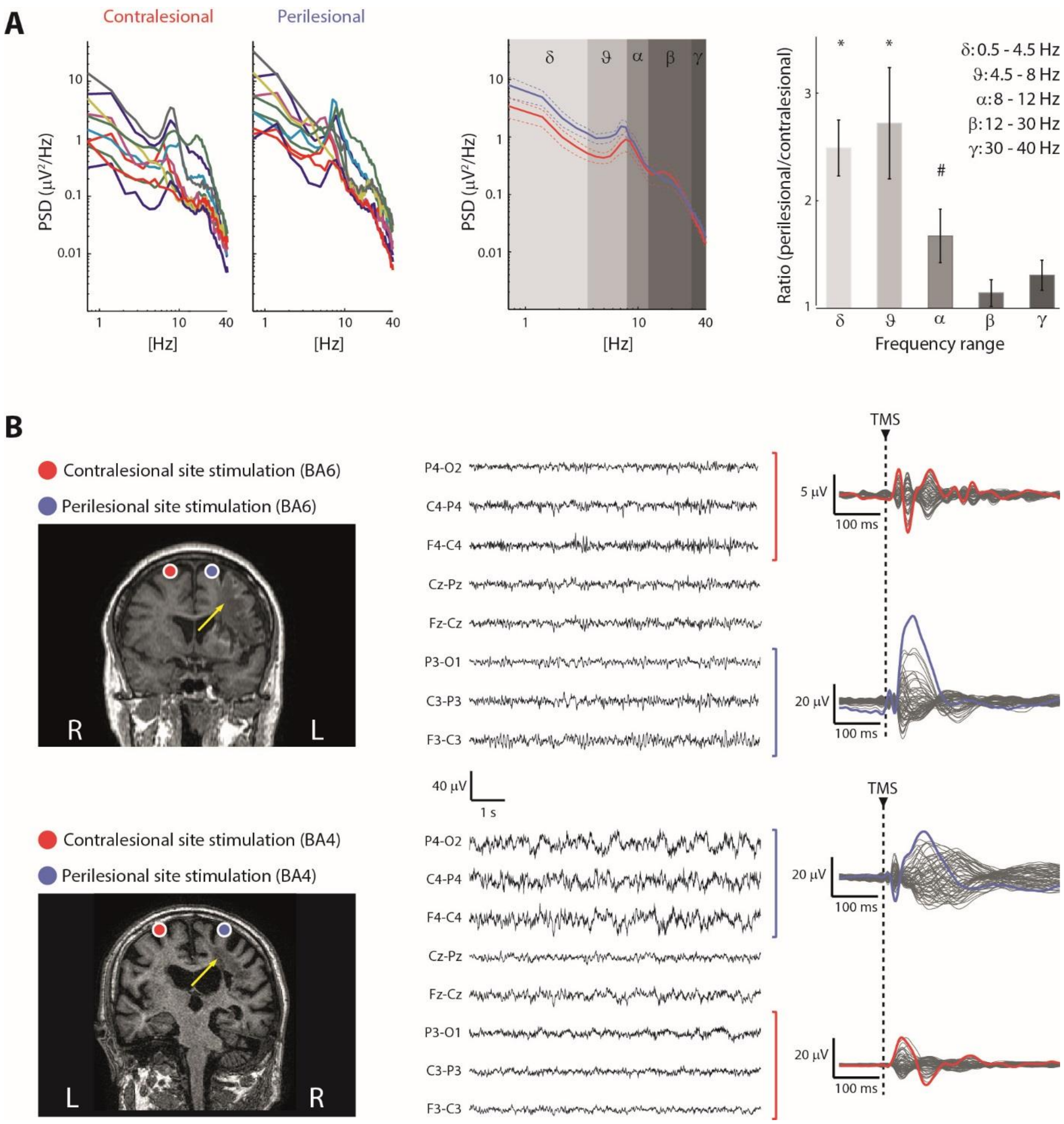

**Fig. S1.** *Resting-state EEG recordings in the MCA ischemia group.* **Panel A.** For each patient, individual Power Spectral Density (PSD) calculated over the same four channels used for TMS/EEG analysis (contralesional, red; perilesional, blue) is shown (left). Average PSD (±SEM) across patients. Shaded gray boxes identify classical frequency ranges (middle). Ratio between perilesional and contralesional site PSD averaged across bins divided into classical frequency ranges: delta (0.5-4.5Hz), theta (4.5-8 Hz), alpha (8-12 Hz), beta (12-30 Hz) and gamma (30-40 Hz) (right). Group analysis showed a significant increase of delta and theta EEG activity over the perilesional site (*: p-values <0.008 for both delta and theta; Bonferroni corrected α: 0.01), confirming the typical EEG pattern found in unilateral brain injuries characterized by cortical infarction *(70)*. Alpha frequency was also found increased over the perilesional site (#: p=0.02 uncorrected). **Panel B.** MRIs and cortical targets as estimated by the Navigated Brain Stimulation system are shown for two representative patients (patient n.4 and n.5 from Table1, left). The yellow arrows highlight lesion location. The visual inspection of EEG (here displayed with a reduced longitudinal bipolar montage focused on the regions explored by TMS, middle) confirmed the PSD findings and highlighted heterogeneous EEG patterns (see also Table S1) characterized by the lateralized presence of either theta rhythms (top) or slow waves (bottom).

Notably, in both cases, applying TMS (dashed vertical line) resulted in a clear-cut evoked EEG slow wave over the perilesional site (blue traces on the right), thus showing the added value of TMS in revealing the presence of perilesional slow waves (and of the associated OFF-periods, not shown here, but see Figure 1, Figure 2 and Figure S2) irrespective of the presence of slow waves spontaneously occurring in the background EEG.

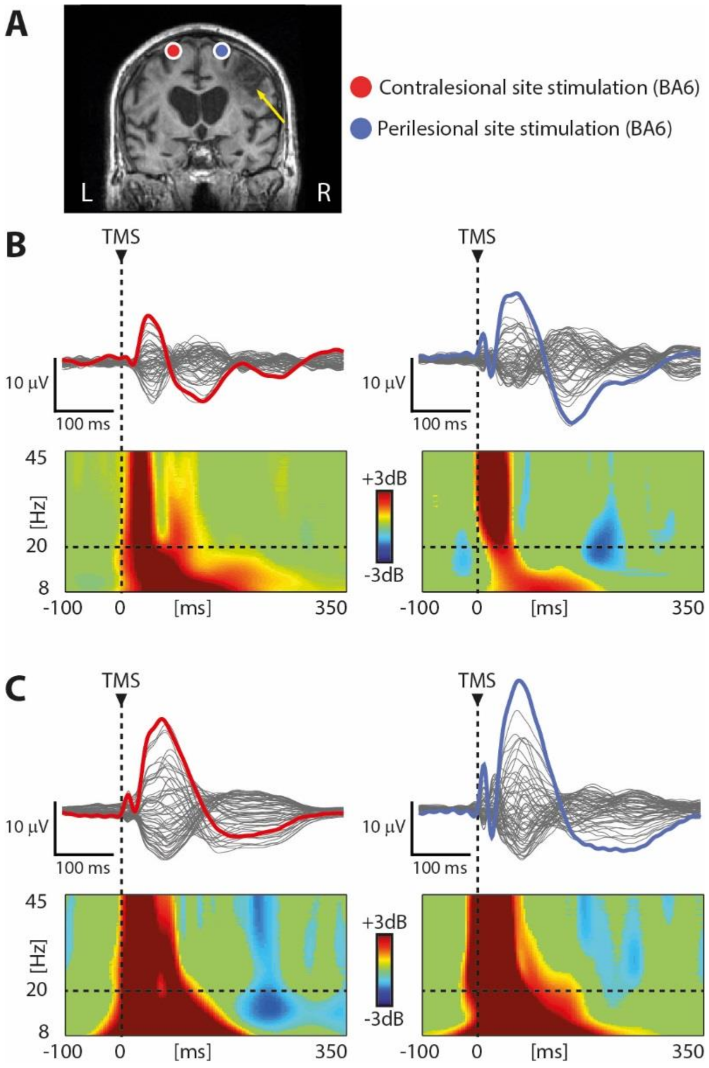

**Fig. S2.** *The TMS-evoked response over the perilesional stimulation site during wakefulness is similar to the typical TMS response ubiquitously observed during NREM sleep.* Results from one representative patient (patient n.14 from Table1) are shown for both the contralesional (red) and perilesional (blue) stimulation sites. **Panel A.** MRI and cortical targets (BA6) as estimated by the Navigated Brain Stimulation system are shown. The yellow arrows highlight lesion location. **Panel B and C.** Butterfly plots of the TMS-evoked EEG potentials recorded at all 60 electrodes (traces) during wakefulness (Panel B) and NREM sleep (Panel C) are depicted. A dashed vertical line marks the occurrence of TMS. Event-related spectral perturbation (ERSP) is presented for the EEG electrode (colored trace) with the largest early response, selected among the four channels closest to TMS. In the ERSP plot, significance for bootstrap statistics is set at $\alpha < 0.05$ (absence of any significant difference form baseline spectrum is colored in green): statistically significant increases of power compared to baseline are colored in red, while blue represents significant power decreases. The dashed horizontal line indicates the 20 Hz frequency bin. Note the similarity between the EEG response and

ERSP features to TMS of the perilesional site stimulation during wakefulness and those observed during sleep over both stimulated sites.

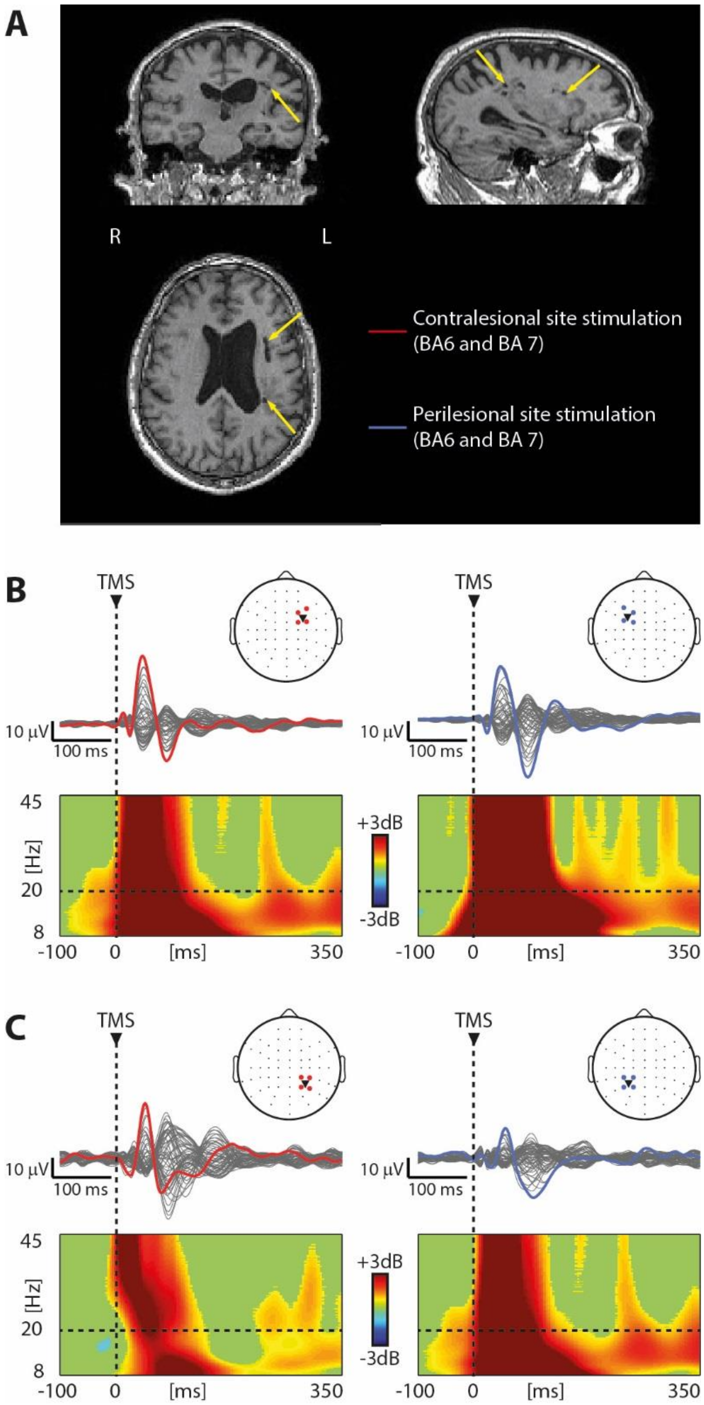

**Fig. S3.** *The absence of TMS-evoked slow waves and OFF-periods irrespective of the stimulated hemisphere and cortical area in the group of patients affected by unilateral lacunar ischemic or haemorrhagic subcortical lesions.* Results from one representative patient (patient n.24 from Table1) are shown for both the contralesional (red) and perilesional (blue) stimulation sites. **Panel A.** MRI and cortical targets (BA6) as estimated by the Navigated Brain Stimulation system are shown. The yellow arrows highlight lesion location. **Panel B and C.** Butterfly plots of the TMS-evoked EEG potentials recorded at all 60 electrodes (traces) during wakefulness (Panel B) and NREM sleep (Panel C) are depicted. A dashed vertical line marks the occurrence of TMS. Event-related spectral perturbation (ERSP) is presented for the EEG electrode (colored trace) with the largest early response, selected among the four channels closest to TMS. In the ERSP plot, significance for bootstrap statistics is set at α < 0.05 (absence of any significant difference form baseline spectrum is colored in green): statistically significant increases of power compared to baseline are colored in red, while blue represents significant power decreases. The dashed horizontal line indicates the 20 Hz frequency bin. Note the absence of TMS-evoked slow waves and OFF-periods across all stimulated sites.

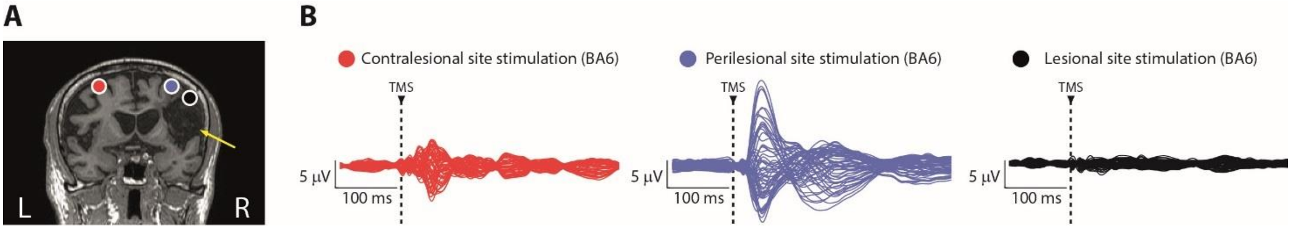

**Fig. S4.** *TMS reveals three distinct cortical reactivity profiles within the same patient.* Results from one representative patient (patient n.10 from Table1, same as in Figure 1) are shown for the contralesional (red), the perilesional (blue) and the lesional (black) stimulation sites. **Panel A.** MRI and cortical targets (BA6) as estimated by the Navigated Brain Stimulation system are shown. The yellow arrow highlights lesion location. **Panel B.** Butterfly plot of the TMS-evoked EEG potentials recorded at all 60 electrodes for the three stimulation sites. Note the absence of EEG reactivity to TMS over the lesion. The dashed vertical line marks the occurrence of TMS.

| Patient number | Lateralized anomalies | Focal anomalies | | | PDR anomalies | | Bilateral anomalies | |
|---|---|---|---|---|---|---|---|---|
| | | **Location** | **Frequency band** | **Incidence** | **Presence** | **Side prevalence** | **Location** | **Side prevalence** |
| **1** | marked | right F-C-T | delta-theta | sub-continuous | yes | right suppressed PDR | N/A | N/A |
| **2** | none | N/A | N/A | N/A | yes | right PDR slowing | N/A | N/A |
| **3** | mild | right T | sharp waves | intermittent | no | N/A | N/A | N/A |
| **4** | mild | left F-C | theta | intermittent | no | N/A | N/A | N/A |
| **5** | marked | right C-P-T | delta | continuous | yes | right suppressed PDR | N/A | N/A |
| **6** | mild | right P | delta | sub-continuous | no | N/A | bifrontal | left |
| **7** | marked | right T-P | delta-theta | sub-continuous | yes | right suppressed PDR | N/A | N/A |
| **8** | none | N/A | N/A | N/A | yes | right suppressed PDR | N/A | N/A |
| **9** | mild | left F-T | delta | sporadic | no | N/A | N/A | N/A |
| **10** | mild | right P-O | theta | sub-continuous | no | N/A | N/A | N/A |

**Table S1.** *Spontaneous EEG clinical assessment of MCA ischemia patients.* F: Frontal. C: Central. T: Temporal. P: Parietal. O: Occipital. PDR: Posterior Dominant Rhythm.

| Patient number | *MCA ischemia* | Patient number | *Severe Multifocal* | Patient number | *Subcortical* |
|---|---|---|---|---|---|
| | **Maximum PCI$^{st}$** | | **Maximum PCI$^{st}$** | | **Maximum PCI$^{st}$** |
| **1** | 54.3 | **11** | 31.8 | **21** | 29.7 |
| **2** | 43.4 | **12** | 41.1 | **22** | 45.7 |
| **3** | 54.3 | **13** | 34.7 | **23** | 44.0 |
| **4** | 33.9 | **14** | 33.7 | **24** | 39.5 |
| **5** | 37.6 | **15** | 26.7 | **25** | 38.7 |
| **6** | 40.9 | **16** | 29.5 | **26** | 53.9 |
| **7** | 33.2 | **17** | *11.7* | **27** | 45.9 |
| **8** | 39.2 | **18** | *12.7* | **28** | 44.0 |
| **9** | 37.5 | **19** | 25.9 | **29** | 42.0 |
| **10** | 36.9 | **20** | 37.1 | **30** | 41.5 |
| **Mean** | 41.1 | **Mean** | 28.5 | **Mean** | 42.5 |
| **SE** | 2.4 | **SE** | 3.1 | **SE** | 2.0 |

**Table S2.** *Maximum global spatiotemporal dynamics of the TMS-evoked responses assessed via the Perturbational Complexity Index (PCI$^{st}$ (27)).* Gray text highlights the two patients (n. 17 and 18) pertaining to the severe multifocal lesions group for which maximum PCI$^{st}$ value was found below the benchmark statistical threshold reported in *(27)*.

| Patient number | Contralesional Natural Frequency (Hz) | Perilesional Natural Frequency (Hz) | Stimulation site |
|---|---|---|---|
| **21** | 13.9 | 12.6 | BA4 |
| **22** | 10.9 | 12.6 | BA4 |
| **23** | 23.5 | 12.3 | BA6 |
| **24** | 14.3 | 11.7 | BA6 |
| **25** | 13.6 | 25.1 | BA6 |
| **26** | 13.8 | 29.5 | BA6 |
| **27** | 10.5 | 10.2 | BA4 |
| **28** | 7.9 | 7.4 | BA4 |
| **29** | 31.2 | 12.9 | BA6 |
| **30** | 18.5 | 18.2 | BA6 |
| **Mean** | 15.8 | 15.3 | |
| **SE** | 2.2 | 2.2 | |

**Table S3.** *Assessment of the frequency of the main oscillatory components of the TMS-evoked response (natural frequency (28)) in the subcortical lesion group.*
